# Phenotype-driven identification of drug targets for post-COVID-19 anosmia

**DOI:** 10.1101/2022.08.03.502673

**Authors:** Victoria M. Catterson, Yolanda Sanchez, Gabriel Musso

**Affiliations:** BioSymetrics Inc

**Keywords:** anosmia, COVID-19, post-COVID sequelea, long COVID, drug target identification

## Abstract

Anosmia (loss of sense of smell) is one symptom of COVID-19 which can linger long after acute infection has passed, with major impact on quality of life. Given the number of people impacted by COVID-19-related anosmia, there is an urgent need to identify effective therapeutics in a faster fashion than using traditional drug discovery and development methods. We used our knowledge graph, the Phenograph, to navigate from phenotypes to genes to drug targets, to rapidly find druggable targets associated with anosmia. This process shortlisted six targets: NRP1, SCN9A, EGR1, VEGFB, PRKCE, and FGFR1. Neuropilin-1 (NRP1) is under active study for its involvement in SARS-CoV-2 infection. Importantly, there is no direct link between anosmia and NRP1 in our knowledge graph; the relationship was inferred through the graph structure. Based on this external validation, we derived hypotheses for the involvement of the remaining five targets in COVID-19-related anosmia, and the mechanism of action desired in a drug candidate to correct the hypothesized dysregulation.

## Introduction

Anosmia (loss of sense of smell) is often considered a milder symptom of COVID-19, and yet it can have a significant impact on quality of life for those affected^1^. Further, smell disruption can linger long after acute infection has passed, with one study reporting 46% of mild COVID-19 cases experiencing some disturbance and 7% total anosmia one year after infection^2^. With no standard treatment, this represents a substantial, current, and growing population of unmet need. While novel therapeutic discovery efforts can take 10 years or more to bring a treatment to the clinic, a target which is known to be druggable (as it is already the target of an approved drug) has the potential to substantially reduce this timeline. For this reason, we consider COVID-19-related anosmia, specifically post-acute infection, to be well suited for target identification through a known-druggable lens.

We used our knowledge graph, the Phenograph (see **Methods**), to identify six druggable targets which could treat COVID-19-related anosmia, by mapping from COVID-implicated phenotypes, to associated genes, and to the products of those genes known to be targeted by one or more drugs. While all six targets have evidence of being druggable, involved in anosmia, and associated with SARS-CoV-2 infection, only one is hypothesized to result in anosmia due to over-expression of the gene product (Early Growth Response 1 (EGR1): critical to the suppression and subsequent reactivation of synaptic activity in the olfactory receptor neurons in the presence of odors). In the drug discovery context,, inhibition is generally easier to address than activation as a mechanism of action, and therefore EGR1 appears to be the strongest candidate for further study.

Our druggable target identification workflow is depicted in Figure 1. The Phenograph listed 51 genes associated with the Human Phenotype Ontology (HPO) term for anosmia, HP:0000458. These genes were ranked and prioritized based on their specificity to anosmia, by counting how many HPO terms (other than those associated with smell and taste) they associated with. That is, a gene linked only to HPO terms for anosmia and decreased taste (hypogeusia, HP:0000224) would rank higher than a gene associated with 100 diverse HPO terms. We performed pathway expansion on the top ranking 10 genes, resulting in 60 candidate targets. These were mapped to drug targets using sources such as the Therapeutic Target Database (TTD), as a proxy for finding druggable targets. Targets were considered druggable if they were associated with at least one drug candidate that advanced to at least a Phase 2 clinical trial, under the assumption that the drugs had passed a Phase 1 safety assessment, but may not have progressed further due to not meeting Phase 2 or 3 efficacy endpoints. (Since such a trial would have been for a different indication, this should not rule out such a target as being druggable.) The final step was to develop a hypothesis of dysregulation in the gene product which would result in anosmia, in order to determine the desirable mechanism of action of a drug candidate to correct post-COVID-19 anosmia.

**Fig. 1.**
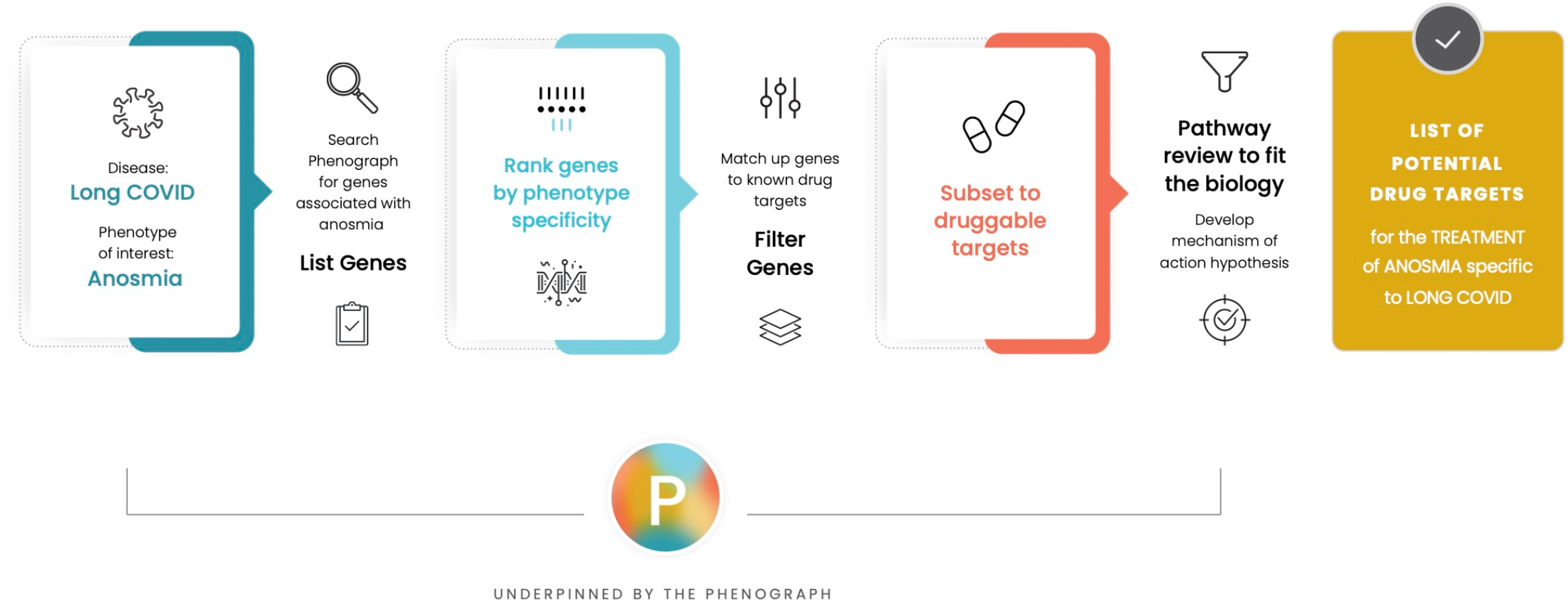
The drug target identification workflow. Each step of the process is underpinned by data extracted from the Phenograph, our knowledge graph which associates diseases, phenotypes, genes, and drugs.

## Results

We found six druggable targets with literature supporting their association with both SARS-CoV-2 infection and anosmia (although none were previously associated specifically with COVID-19 related anosmia) (Table 1). All six targets have evidence of being druggable, as indicated by Phase 2 or later-stage clinical trials for drug candidates, although for different indications. Two of the targets have drugs either in trials or tested in pre-clinical models for SARS-CoV-2 efficacy (NRP1 and FGFR1). While we can suggest a role for all six targets in anosmia, only one has a hypothesized over-expression dysregulation (EGR1). This clear directionality of expression in disease prompted a focus on EGR1 as an attractive target for further drug development.

**Table 1:** Druggable targets and hypothesized anosmia-causing dysregulation found using the Phenograph knowledge graph

| Target | Hypothesized dysregulation leading to anosmia |
| --- | --- |
| **Neuropilin-1 (NRP1)** | Under-expression |
| **Sodium Voltage-Gated Channel Alpha Subunit 9 (SCN9A)** | Under-expression |
| **Early Growth Response 1 (EGR1)** | Over-expression |
| **Vascular Endothelial Growth Factor B (VEGFB)** | Under-expression |
| **Protein Kinase C Epsilon (PRKCE)** | Under-expression |
| **Fibroblast Growth Factor Receptor 1 (FGFR1)** | Under-expression |

## Discussion

While the Phenograph associates phenotypes to genes to drugs, we tried to determine a feasible hypothesis for each target, and what mechanism of action for a drug candidate would correct dysregulation. The sections below elaborate further on each target, exploring the available evidence for the association with SARS-CoV-2 infection and anosmia, and discussing the potential mechanism of action.

### Neuropilin-1 (NRP1)

Our hypothesis is that after infection by SARS-CoV-2, olfactory bulb (OB) epithelial damage is repaired with inefficient olfactory nerve axon targeting, due to NRP1 dysregulation. In healthy subjects, the olfactory nerve axon is directed to attach to the correct zone of the OB where NRP1 is highly expressed. Low availability of NRP1 in the presence of binding to SARS-CoV-2 results in poor targeting, and therefore odor-generated action potentials cannot be transmitted to the olfactory nerve, resulting in anosmia.

In support of this hypothesis, NRP1 was identified as a co-receptor that significantly potentiates SARS-CoV-2 infectivity^3,4^. Monoclonal antibodies, siRNA, and/or inhibitors targeting NRP1, were found to significantly block viral entry into cells, demonstrating a role for NRP1 in infection by this virus^3^. Further, NRP1 binds furin-cleaved substrates, and a SARS-CoV-2 mutant with an altered furin cleavage site did not depend on NRP1 for infectivity^4^. Finally, analysis of olfactory epithelium obtained from human COVID-19 autopsies revealed SARS-CoV-2 infected NRP1-positive cells facing the nasal cavity^3^. Altogether, these results provide insight into SARS-CoV-2 cell infectivity and demonstrate that NRP1 serves as a host factor for the SARS-CoV-2 virus, thus defining a new potential target for antiviral intervention.

In relation to anosmia, levels of NRP1 are directly involved in guiding olfactory axons to the correct target zone of the olfactory bulb (OB)^5^. One study proposed that the inverse link between COVID-19 severity and anosmia can be explained by NRP1-mediated cell entry to the sensory neurons for smell^6^. A post mortem study of COVID-19 patients showed that non-neuronal (epithelial) cells are preferentially targeted by SARS-CoV-2 in the OB^7^. Therefore, our hypothesis would explain how non-neuronal targeting can suppress neuronal signaling.

In addition, VEGFA/NRP1 signaling has been implicated in nociceptor activity and neuropathic pain^8^. The SARS-CoV-2 spike protein subverts VEGFA/NRP1 pronociceptive signaling, hijacking NRP1 signaling to ameliorate VEGFA-mediated pain. This raises the possibility that pain, as an early symptom of COVID-19 disease, may be directly dampened by the SARS-CoV-2 spike protein.

#### Mechanism of action for NRP1

The literature evidence discussed above shows we are not the first to implicate NRP1 for its role in COVID-19. Molecular docking studies have proposed structures for compounds predicted and then validated to bind to NRP1^9,10^, including one drug which is currently undergoing clinical trials.

This drug is a small molecule previously approved for pancreatitis. The literature supporting its COVID-19 trial suggests it directly targets SARS-CoV-2, but it is also predicted to bind to NRP1^10^. With this mechanism, it offers the potential to prevent or reduce the severity of SARS-CoV-2 infection, and hence avoid anosmia as a secondary effect. However, to address anosmia after acute infection has passed, the required mechanism is an activator of NRP1 rather than an inhibitor.

### Sodium Voltage-Gated Channel Alpha Subunit 9 (SCN9A)

Multiple Transient Receptor Potential (TRP) channels are implicated in viral entry via the ACE2 receptor (the primary route for SARS-CoV-2 infection), leading to an influx of intracellular Ca+2 ions^11^. These Ca+2 ions provoke an inflammation reaction through the Raf/MEK/ERK pathway^11^. A drop in extracellular pH from the Ca+2 ions causes the Acid Sensing Ion Channel Subunit 1 channel (ASIC1: a proton-gated sodium channel) to open, leading to Na+ ion influx^11^. We posit that this Na+ influx leads to dysregulation of the SCN channels, and hence to anosmia, as the dysregulation of the voltage-gated sodium channel NaV1.7 encoded by *SCN9A* has been shown to prevent sensory signals passing through the olfactory nerve^12^.

Further support for this hypothesis comes from studies showing that humans with a loss of function (LoF) mutation in *SCN9A* have an insensitivity to pain^13^, while gain of function (GoF) mutations have been associated with pain disorders^14^. LoF mutation in *SCN9A* in mice was shown to give rise to anosmia due to electrical signals failing to transmit through the first synapse of the olfactory nerve^14^, and specifically due to inhibition of neurotransmitter function^12^. Hence, dysregulation of the NaV1.7 channel due to the immune response to ACE2 receptor infection by SARS-CoV-2 could have the same anosmic effect as LoF mutations in *SCN9A*.

#### Mechanism of action for SCN9A

There has been substantial drug discovery effort targeting *SCN9A*, due primarily to the analgesic effect of a LoF mutation in this gene, but such efforts focus on developing sodium channel blockers with the aim of reducing signaling and therefore ameliorating pain. To address anosmia, the goal would be to enhance neurotransmitter activity, not to block it. Therefore, the desired mechanism of action is an activator of SCN9A.

### Early Growth Response 1 (EGR1)

Our hypothesis is that overexpression of *EGR1* in the OB resulting from SARS-CoV-2 infection leads to a continuously elevated level of tyrosine hydroxylase (*Th*) in the olfactory receptor neurons, thus inhibiting synaptic activity and preventing the signal from an odor being transmitted.

Under normal conditions, *EGR1* is expressed at low levels in the OB, but is transiently highly expressed in the presence of odors^15^. EGR1 regulates transcription of *Th*, which is expressed in the interneurons of the OB in response to an odor, with the effect of rate limiting production of dopamine and thus inhibiting synaptic activity between multiple OB neurons, including olfactory receptor, mitral, and tufted neurons^15^. Over-expression of EGR1 can therefore be expected to continuously suppress such synaptic activity. Some evidence shows that antisense mutations in EGR1 in fruit bats prevent odor memory formation^16^, potentially due to anosmia.

The evidence for SARS-CoV-2 association with EGR1 is more circumstantial than for the previous targets, but does support there being an effect. A lung transcriptome study showed that the activity of EGR1 as a transcription factor was modulated by viral proteins, with the effect of inhibiting hypoxia response^17^, a dysregulation which had been seen prior to COVID-19^18^. Secondly, expression profiling of sensory neurons after SARS-CoV-2 infection showed statistically significant increase in *EGR1* expression^19^. Given that NRP1 is a secondary route for viral entry of SARS-CoV-2, and NRP1 is highly expressed in the OB, olfactory sensory neurons could be infected by this route with corresponding increased expression of EGR1 as a result.

#### Mechanism of action for EGR1

Since over-expression of EGR1 is hypothesized to result in anosmia, a candidate drug should be an inhibitor of EGR1, highlighting EGR1 as an attractive target for future drug development efforts.

### Vascular Endothelial Growth Factor B (VEGFB)

The hypothesis for VEGFB is more circumstantial than for other targets, but derives strength through association with NRP1. VEGFB binds to NRP1 and vascular endothelial growth factor receptor 1 (VEGFR1) seemingly to activate downstream signalling pathways important for skeletal muscle growth and repair^20^. However, being a homolog of VEGFA, it is thought that VEGFB may offer similar or redundant functionality to VEGFA^20^. As such, the known SARS-CoV-2 spike protein subversion of the VEGFA/NRP1 pronociceptive signaling^8^ may also apply here. However, it should be noted that the biological function of VEGFB remains “enigmatic” as of 2021^20^, hence its role in anosmia is implicated through association with NRP1, and the role of NRP1 in guiding olfactory axons to the OB.

#### Mechanism of action for VEGFB

We hypothesize above that an activator of NRP1 would address COVID-related anosmia by directing axons to insert to the correct location in the OB. Following similar logic, if VEGFB/NRP1 is involved in sensory signaling, activation of VEGFB is required to enhance signaling.

### Protein Kinase C Epsilon (PRKCE)

The support for PRKCE in anosmia is based on its involvement in neuronal signaling, its high expression levels in the OB, the involvement of various members of the protein kinase C (PKC) family in signaling triggered by SARS-CoV-2 infection^21^, and increased expression of PKC members in a cell line resistant to viral infection^22^.

While not known to associate with anosmia, PRKCE expression is third highest in OB tissue, after cerebral cortex and lung, according to the FANTOM5 dataset^23^. Its role in neuronal signaling has been well explored in relation to bipolar disorder, with specific work showing increased PKC activity in platelets and post-mortem brains of patients with bipolar disorder^24^, and by implication through the effects of drugs lithium and valproate being to reduce PRKCE and PRKCA selectively^25^.

While bipolar disorder implicates overexpression of PRKCE leading to dysregulated neurotransmitter release and over-excitation of neurons (hence addressed by drugs such as lithium which suppress PRKCE), our hypothesized role in anosmia is an under-expression of PRKCE leading to failure of neuronal signaling in the presence of odors.

#### Mechanism of action for PRKCE

Based on the current hypothesis, a candidate drug would be required to activate PRKCE to address the under-expression underlying anosmia through suppression of neuronal signaling.

### Fibroblast Growth Factor Receptor 1 (FGFR1)

Mutation of FGFR1 is a known cause of Kallmann Syndrome, a congenital condition characterized by idiopathic hypogonadotropic hypogonadism and anosmia^26^. Anosmia due to Kallmann Syndrome is caused by the olfactory epithelium failing to develop fully or at all, leading to an under-developed OB^27^.

While not directly implicated in SARS-CoV-2 infection, FGFR1 is a cofactor of infection with viruses such as influenza^28^, and FGFR1 signalling was shown to explain the association of Epstein-Barr virus with nasopharyngeal cancer^28^. This weaker evidence of a role for FGFR1 in viral infection is bolstered by the strong association with causing anosmia, and hence our hypothesis is that dysregulation of FGFR1 due to infection results in failure to adequately repair olfactory epithelium after acute COVID-19, and hence anosmia is prolonged past the acute infection.

#### Mechanism of action for FGFR1

Since LoF mutation of FGFR1 is known to cause anosmia, we hypothesize that SARS-CoV-2 infection is causing anosmia through under-expression of FGFR1. Hence, the desired mechanism of action for a drug candidate is an activator of FGFR1.

Notably, one study recently screened a number of drugs *in vitro* against human coronaviruses, including an inhibitor of FGFR1-3 approved for various lung disease indications such as idiopathic pulmonary fibrosis, which showed only “moderate” inhibition of SARS-CoV-2 cytotoxicity^29^. However, we do not draw conclusions either way from this result, as the mechanism of action to address acute infection may be different from the mechanism of action needed to address post-acute symptoms (as in the case of NRP1, above). Rather, we find it notable that others have explored this target and even tested a drug with this target for SARS-CoV-2 infection more broadly.

## Conclusions

We used our Phenograph knowledge graph to traverse from phenotype to genes to drugs, with the aim of identifying druggable targets which may resolve post-COVID-19 anosmia. This method uncovered six such targets with literature support for association with both SARS-CoV-2 infection and anosmia. When considering the mechanism of action of appropriate drug candidates, one target has a hypothesized over-expression of the gene product as the cause of anosmia, providing clear directionality to the required mechanism. Inhibition of a target is generally considered as more accessible from the perspective of characterizing novel drug-like molecules, as the dynamic range and compound efficacy in biochemical and/or cellular assays used in early drug discovery can be readily ascertained. Overall, these considerations make EGR1 an attractive target for further study.

These results highlight the applicability of curated knowledge networks in identifying avenues for druggable target identification and formulating new testable hypotheses, particularly for newly-identified disorders where there is a requirement for expedited target nomination.

## Materials and Methods

The Phenograph is built as a knowledge graph drawing on data from public and proprietary sources. It includes human disease, phenotype, and gene information, as well as gene and phenotype data for model organisms such as mouse and zebrafish. Data is integrated from public sources such as Open Targets, BioMart, the Human Phenotype Ontology, the Jackson Laboratory, LOINC, and ZFIN; and an image of this version of the Phenograph has been cached to facilitate reproducibility of the presented analysis. It currently contains over 250k nodes and 11.3M edges, some contributed by a process of machine-learning-based gene-phenotype prediction. All of the human disease biology used for this study is curated from public sources. However, the ease of navigation from phenotype to gene to drug, and the ability to collect and rank a list of genes within the Phenograph saved considerable time and effort over performing this same work directly from those sources.

To arrive at the final list of 6 targets and potential drugs, the process involved gene selection, gene ranking, pathway expansion, drug matching, and final investigation, detailed below and depicted in **Figure 1**.

Genes were selected based on any known association with anosmia. Data such as the Human Phenotype Ontology’s Phenotype2gene dataset^30^ annotates HPO terms with genes based on evidence of genetic association studies. This and similar data within the Phenograph was searched for the HPO term HP:0000458 (anosmia), and all resulting genes added to our list. This gave a list of 51 genes.

Gene ranking then prioritized this list of 51 according to a score of how specific the gene is to anosmia. By performing the reverse search of gene to phenotype for each gene in the list, we could count how many HPO terms a gene is known to associate with. We created a shortlist of anosmia-related phenotypes and removed them from the results, on the assumption that these terms do not indicate a lack of specificity to anosmia. This list was:

- Abnormality of taste sensation HP:0000223
- Hypogeusia HP:0000224 (reduced taste)
- Ageusia HP:0041051 (loss of taste)
- Abnormality of the sense of smell HP:0004408
- Hyposmia HP:0004409 (decreased perception of odours)
- Anosmia HP:0000458

That is, a gene’s score was the number of HPO terms not on this list that a gene is known to associate with. Genes were prioritized by this score, where smaller is better. The 10 genes with the lowest scores were taken as a shortlist.

Pathway expansion was performed to increase those 10 genes to 60. The assumption here is that a specific gene may not be druggable, but targeting an alternative in the same pathway could have the desired effect. Data on pathways was drawn from GeneMANIA^31^, which aggregates 6 different sources from Pathway Commons and others.

Druggable target filtering drew on data from the Therapeutic Target Database (TTD)^32^. We considered only drugs at the stage of Phase 2 clinical trial or further, with the assumption that drugs which have failed a Phase 2 trial for a different indication may still target the gene product of interest, but have also passed the safety assessment of a Phase 1 trial. The targets of all such drugs were considered “druggable targets”. The 60 genes on our pathway-expanded list were searched within this list, which automatically dropped the number to 12 druggable targets.

The final investigation is where we reviewed literature on the target to find evidence of a link with SARS-CoV-2 infection in addition to anosmia. Those targets with papers supporting evidence for both were kept on the list, and a hypothesis developed for the required mechanism of action of a candidate drug. This step is the only manual part of the process, and it had to be performed manually due to the relative speed of knowledge generation about SARS-CoV-2 and COVID-19. Associations between genes, drugs, and SARS-CoV-2 have not yet entered the public datasets from which the Phenograph draws the majority of its human disease knowledge. This final investigation into disease biology reduced the number of targets to 6, as presented in Table 1.

## Acknowledgments

With thanks to Nishanth Merwin, Sidney Elmer, Shahrzad Hosseini, Kevin Ha, Simon Eng, William Stanford, and Stacie Calad-Thomson for discussions and feedback on the workflow; and to Steven Bishop for software design and development of the Phenograph.

